# A novel function of sphingosine kinase 2 in the metabolism of sphinga-4,14-diene lipids

**DOI:** 10.1101/2020.02.14.949974

**Authors:** Timothy A Couttas, Yepy H Rustam, Huitong Song, Yanfei Qi, Jonathan D Teo, Jinbiao Chen, Gavin E Reid, Anthony S Don

**Author notes:** Corresponding author: Anthony S Don, Centenary Institute and NHMRC Clinical Trials Centre, Faculty of Medicine and Health, The University of Sydney, Camperdown, NSW, 2006, Australia.

## Abstract

The number, position, and configuration of double bonds in lipid acyl chains affects membrane packing, fluidity, and recruitment of signalling proteins. Studies on mammalian sphingolipids have focused on those with a saturated sphinganine or mono-unsaturated sphingosine long chain base. Sphingolipids with a diunsaturated sphingadiene base have been reported but are poorly characterised. Employing high-resolution untargeted mass spectrometry, we observed marked accumulation of lipids containing a sphingadiene base, but not those with a more common sphingosine backbone, in the hippocampus of mice lacking the metabolic enzyme sphingosine kinase 2 (SphK2). Applying ultraviolet photodissociation tandem mass spectrometry (UVPD-MS/MS) the double bonds were confidently assigned to the C4-C5 and C14-C15 positions of the sphingoid base. Sphingosine kinases are involved in lysosomal catabolism of all sphingolipids, producing sphingoid base phosphates that are irreversibly degraded by sphingosine 1-phosphate lyase. Both SphK1 and SphK2 phosphorylated sphinga-4,14-diene as efficiently as sphingosine, however deuterated tracer experiments demonstrated that ceramides with a sphingosine base are more rapidly metabolised in cultured cells than those with a sphingadiene base. SphK2 silencing significantly impeded the catabolism of both sphingosine- and sphingadiene-based sphingolipids. Since SphK2 is the dominant sphingosine kinase in brain, we propose that accumulation of sphingadiene lipids in SphK2-deficient brains results from the intrinsically slower catabolism of sphingadiene lipids, combined with a bottleneck in the catabolic pathway created by the absence of SphK2. We speculate that accumulation of these lipids in the absence of SphK2 function may affect the fluidity and signalling properties of cell membranes.

## Introduction

Sphingolipids are a large and diverse family of lipids, heavily enriched in the mammalian brain. They are important regulators of cell signaling and physiology in eukaryotes (1, 2). The common constituent of sphingolipids is their sphingoid base, which in mammals is predominantly 18-carbon sphingosine that contains a double bond at the C4-5 position (d18:1) (Figure 1a). Lipidomic studies often also quantify sphingolipids with the less abundant saturated dihydrosphingosine (d18:0) backbone (2).

**Figure 1.**
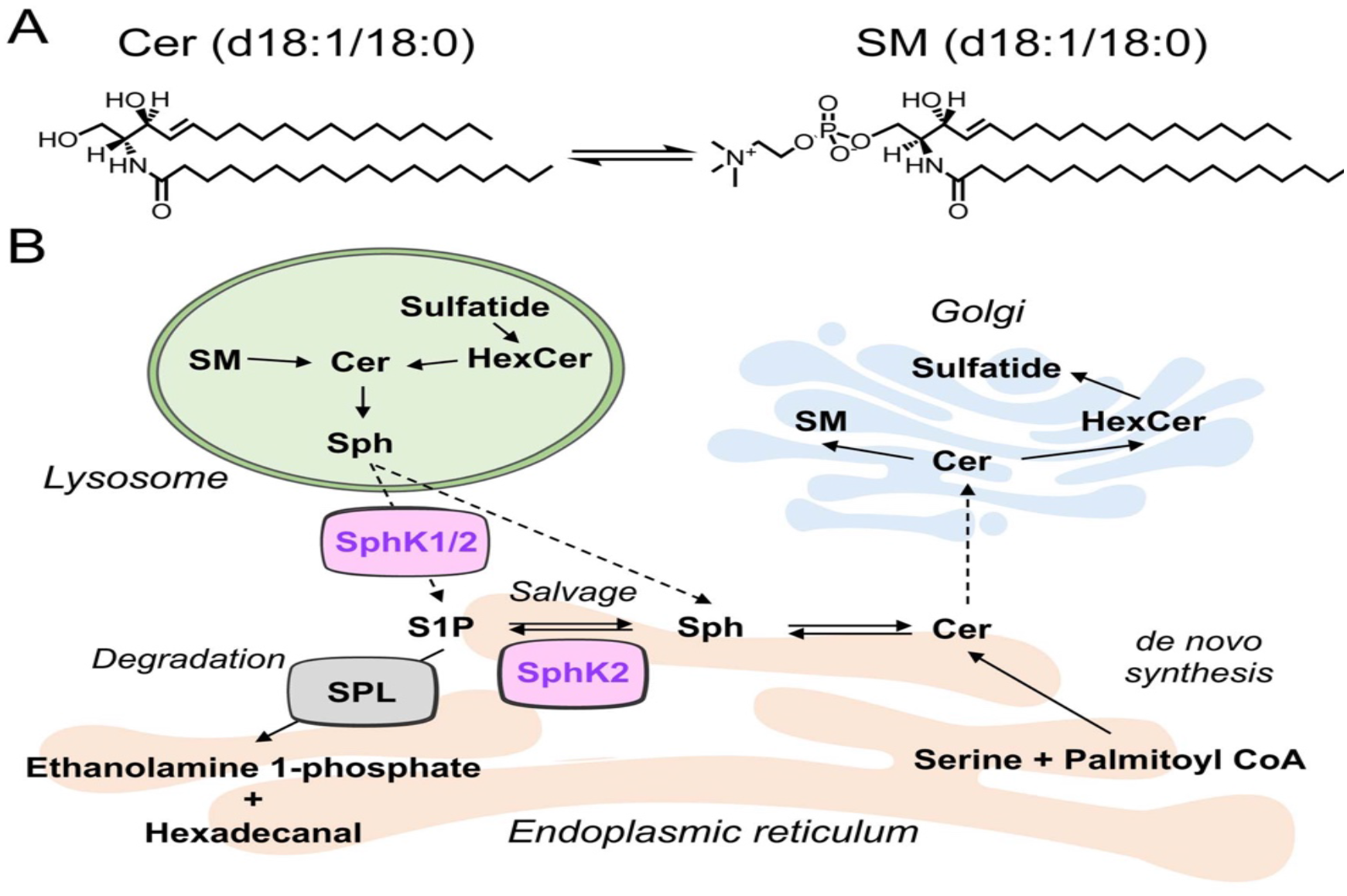
Sphingolipid structure and basic metabolism. (A) Structure of d18:1/18:0 ceramide and SM. (B) Sphingolipid biosynthesis and breakdown. *De novo* synthesis of sphingolipids occurs in the endoplasmic reticulum, producing ceramides from serine and palmitoyl CoA. Ceramide transfer protein (CERT) transfers ceramides to the Golgi, where they are converted to SM and glycosphingolipids such as HexCer and sulfatide. Sphingolipid breakdown in lysosomes results in the generation of sphingosine (Sph), which can be recycled back to ceramide in the salvage pathway or phosphorylated by SphK1 or SphK2, forming S1P. S1P can also be reutilised for sphingolipid synthesis in the salvage pathway or broken down in the endoplasmic reticulum by S1P lyase (SPL).

The core diacyl sphingolipid structure is ceramide, which contains a sphingoid base amide-linked to a fatty-acyl chain (Figure 1A). These fatty-acyl chains vary from 14 to 26 carbons in length, and are generally saturated or mono-unsaturated. Enzymatic addition of different headgroups to the primary hydroxyl of ceramide yields the different classes of sphingolipids, which include sphingomyelin (SM), hexosylceramide (HexCer) and sulfatide.

The length and degree of saturation of the sphingoid base and amide-linked acyl chains of sphingolipids can have a marked impact on their physiological function. For instance, d18:1/16:0 ceramide (C16 ceramide) antagonizes insulin signalling and promotes physiological insulin resistance, whereas its saturated counterpart C16 dihydroceramide (d18:0/16:0 ceramide), which lacks the C4–5 double bond in the sphingoid base, does not (3). In direct contrast to C16 ceramide, C24 ceramides (d18:1/24:0 and d18:1/24:1) protect against the development of insulin resistance in the liver (4, 5). The number, position and configuration (cis or trans) of double bonds on both the sphingoid base and fatty-acyl chain greatly affects the membrane packing properties of sphingolipids, which can in turn affect their cellular signaling functions (6, 7).

Tandem mass spectrometry with conventional collision-induced dissociation (CID) allows for characterisation of lipid headgroups, acyl chain length, and number of carbon-carbon double bonds, but provides no information on the position of the double bonds. UVPD-MS/MS is an emerging approach for structural characterisation of lipids in which high-energy excitation through absorption of 193 nm UV photons causes direct dissociation, yielding product ion and structural data that is not observed with conventional CID (8–12). Importantly, 193 nm UV excitation results in characteristic dissociation of bonds adjacent to carbon-carbon double bonds in lipid acyl chains, allowing the location of double bonds within the acyl chains to be determined.

Phosphorylation of sphingosine and other sphingoid bases by sphingosine kinases 1 and 2 (SphK1 and 2) forms the lipid signalling molecule sphingosine 1-phosphate (S1P) and related sphingoid base phosphates (13, 14). S1P executes a wide array of cell signalling roles both through a family of five G-protein coupled receptors that are specific to S1P (13, 15), and through specific intracellular S1P binding targets such as histone deacetylases (16). In addition to forming the signalling molecule S1P, sphingosine phosphorylation is an essential and evolutionarily-conserved step in the catabolism of all sphingolipids. This proceeds through irreversible degradation of S1P by S1P lyase, forming hexadecenal and ethanolamine phosphate (Figure 1B) (1, 2).

As the only pathway for complete catabolism of sphingolipids, sphingosine kinases and S1P lyase are key regulators of sphingolipid homeostasis (17). In mice, loss of S1P lyase is lethal within a few weeks of weaning, attributed to a range of gross developmental defects (18, 19) and pronounced deregulation of lipid levels in multiple organs (20). The absence of S1P lyase causes increased sphingolipid biosynthesis via the salvage pathway, which is associated with reduced *de novo* sphingolipid biosynthesis, thus causing widespread disruption to lipid metabolism (21). Loss of both SphK1 and SphK2 is also lethal, however the mice die with defects in vasculogenesis and neurogenesis attributed to loss of S1P signalling functions (22). Mice lacking either SphK1 or SphK2 exhibit no obvious phenotypic abnormalities.

We have previously reported loss of S1P and SphK2 activity early in the pathogenesis of Alzheimer’s disease (23), and recently demonstrated that loss of SphK2 sensitizes to hippocampal atrophy and myelin loss in a mouse model of Alzheimer’s disease (24). We have also shown heightened anxiety in mice lacking SphK2 (25). SphK2 catalyses the majority of S1P synthesis in the brain (25), and an important role for SphK2 in protection against brain atrophy following cerebral ischaemia has been established (26). Here, we report pronounced and selective accumulation of sphingolipids containing a second double bond in the sphingoid base, termed sphingadienes, in the hippocampus of SphK2 knockout (SphK2^−/−^) mice. Employing the emerging technique of UVPD-MS/MS, the double bonds were unambiguously assigned to positions C4-C5 and C14-C15 in the sphingoid base backbone of these lipids. In metabolic tracing experiments, we show that these sphingadiene-containing lipids are catabolized more slowly than the more abundant sphingosine-based sphingolipids, and therefore propose that sphingadiene lipids accumulate in brains of SphK2^−/−^ mice due to poor degradation via the sphingosine kinase-S1P lyase pathway.

## Materials and Methods

### Lipid standards

All lipids, with the exception of 17:0/17:0/17:0 triacylglycerol (TAG) (Cayman Chemical, USA), were purchased from Avanti Polar Lipids (distributed by Sigma Aldrich, Castle Hill, NSW, Australia). All lipid extractions were spiked with an internal standard mixture comprising of 0.2 nmole of d17:1 Sph (860640) and S1P (860641), and 1 nmole of each of the following: d18:1/12:0 SM (860583), d18:1/17:0 ceramide (860517), 18:1/12:0 glucosylceramide (860543), d18:1/12:0 sulfatide (860573), 19:0/19:0 phosphatidylcholine (850367), 17:0/17:0 phosphatidylethanolamine (830756), 17:0/17:0 phosphatidylserine (840028), 17:0/17:0 d5-diacylglycerol (110580), 17:0/17:0 phosphatidylglycerol (830456) and 17:0/17:0/17:0 TAG (16489). Deuterated (D7) sphingosine (860657), d18:1 sphingosine (860490), and d18:2 sphinga-4,14-diene (860665) were purchased from Avanti Polar Lipids.

### Mouse tissue samples

Mice lacking functional SphK1 (SphK1^−/−^) or SphK2 (SphK2^−/−^) were derived by breeding heterozygous parents (i.e. SphK1^+/−^ × SphK1^+/−^ or SphK2^+/−^ × SphK2^+/−^), allowing us to compare SphK1^−/−^ and SphK2^−/−^ mice to their wild-type (WT) littermates (i.e. SphK1^+/+^ or SphK2^+/+^). Genotyping was performed as described previously (22). Mice were housed in ventilated cages with enrichment consisting of nesting material and a plastic dome, and were given access to food and water *ad libitum*. For untargeted mass spectrometry analysis, we used groups of 6 male SphK2^−/−^ and WT mice at 8 months of age, or 6 (3 male and 3 female) SphK1^−/−^ and 8 (3 male and 5 female) WT mice at 6 months of age. A separate cohort of male SphK2^−/−^ (n = 9) and WT (n = 11) mice, aged 6 months, were used for targeted lipidomic analysis of the hippocampus. Research was approved by the Animal Care and Ethics Committee of the University of Sydney (2017/1284) and Sydney Local Hospital District Animal Welfare Committee (2014/007), in accordance with the Australian Code of Practice for the Care and Use of Animals for Scientific Purposes.

### Cell culture

The MO3.13 oligodendrocyte cell line was kindly provided by Prof Brett Garner, University of Wollongong, and cultured in DMEM medium supplemented with 10% fetal bovine serum and 2 mM L-glutamine. Cell culture reagents were purchased from Life Technologies.

#### Stable SphK2 knock-down

Short hairpin RNA (shRNA) sequences targeting SphK2 (A: CCGGCTACTTCTGCATCTACACCTACTCGAGTAGGTGTAGATGCAGAAGTAGTTTTTG; and B: CCGGCAGGATTGCGCTCGCTTTCATCTCGAGATGAAAGCGAGCGCAATCCTGTTTTTG) and lentiviral scramble control shRNA, in MISSION® pLKO.1-puro Lentiviral vector (#SHC001), were purchased from Sigma Aldrich. The lentivirus was produced by co-transfection of HEK293T cells with pMD2.G (#12259), pMDLg/pRRE (#12251) and pRSV-Rev (#12253) viral packaging and envelope plasmids, supplied by Addgene (deposited by Dr Didier Trono) (27). Lentiviral supernatants were added to cultured MO3.13 cells for 72 h in the presence of 8 μg/ml polybrene (Sigma, #H9268). Stably-transfected cells were selected with 2.5 μg/mL puromycin (ThermoFisher Scientific, #A1113803) for 12 days prior to use as a cell pool.

#### Metabolic labelling with D7-sphingosine

MO3.13 cells were grown to 80% confluency in 6-well plates, cultured overnight in serum-free OptiMEM I culture medium (#31985062, Life Technologies) supplemented with 2 mM L-glutamine, then incubated with 1 μM D7-sphingosine for 2 h. The medium was then replaced with fresh OptiMEM I, and cells were incubated in quadruplicate for a further 0, 30, 60 or 120 min before washing once with PBS, then lysing in 250 μL of 20 mM Hepes, pH 7.4, 10 mM KCl, 1 mM dithiothreitol, 3 mM β-glycerophosphate and cOmplete™ EDTA-free protease inhibitor cocktail (Sigma Aldrich). Cells were lysed and homogenised by disruption in a Bioruptor Q800R2 Sonicator (QSonica). Protein concentration was determined with Pierce BCA assay (ThermoFisher Scientific). A total of 200 μL lysate protein was used for lipid extraction and sphingolipid quantification by targeted LC-MS/MS. Results were normalised for protein concentration. Experiments were performed on two stably transfected shSphK2 cell lines: SphK2A and SphK2B, compared to cells transfected with the control shRNA.

### Lipid extraction

Lipids were extracted from dissected hippocampus, liver tissue and cell extracts tissue using a two-phase procedure with methyl-tert-butyl ether (MTBE)/methanol/water (10:3:2.5, v/v/v) (28, 29). Hippocampus (10-20 mg) and liver tissue (5-10 mg) were pulverised over dry ice, and transferred to 10 mL glass tubes containing 1.7 mL of MTBE, 500 μl of methanol, and 0.01% butylated hydroxytoluene, spiked with the internal standard mixture. Samples were sonicated in an ice-cold sonicating water bath (Unisonics Australia) for 3 × 30 min. Phase separation was induced by addition of 417 μL of mass spectrometry-grade water, followed by vortexing and centrifugation at 1000 × g for 10 min. The upper organic phase was collected in 5 mL glass tubes. The lower phase was re-extracted by adding 1 mL MTBE, 300 μL methanol, and 250 μL water. The organic phases were combined and dried under vacuum in a Savant SC210 SpeedVac (ThermoFisher Scientific), reconstituted in 300 μL methanol and stored at −20 °C for subsequent analysis.

### Lipid Quantification by LC-MS/MS

#### Untargeted Lipidomics

Lipid extracts from hippocampus tissue were analysed on a Q-Exactive HF-X mass spectrometer with heated electrospray ionization (HESI) probe and Vanquish UHPLC system (Thermo Fisher, Breman, Germany). Extracts were resolved on a 2.1 × 100 mm Waters Acquity C18 UPLC column (1.7 μm pore size), using a 25 min binary gradient at a 0.28 mL/min flow rate. HPLC gradient was as follows: 0 min, 70:30 A/B; 3 min, 70:30 A/B; 4.5 min, 57:43 A/B; 5.5 min, 45:55 A/B; 8 min, 35:65 A/B; 13 min, 15:85 A/B; 14 min, 0:100 A/B; 20 min, 0:100 A/B; 20.2 min, 70:30 A/B and 25 min, 70:30 A/B. Solvent A: 10 mM ammonium formate, 0.1% formic acid in acetonitrile:water (60:40); Solvent B: 10 mM ammonium formate, 0.1% formic acid in isopropanol:acetonitrile (90:10). Data was acquired in full scan/data-dependent MS2 (full scan resolution 70,000 FWHM, scan range 400–1200 *m*/*z*) in both positive and negative ionization modes. The ten most abundant ions in each cycle were subjected to MS2, with an isolation window of 1.4 *m*/*z*, collision energy 30 eV, resolution 17,500 FWHM, maximum integration time 110 ms and dynamic exclusion window 10 s. An exclusion list of background ions was used based on a solvent blank. An inclusion list of the [M+H]^+^ and [M-H]^−^ ions was used for all internal standards. LipidSearch software v4.1.30 (Thermo Fisher) was used for lipid annotation, chromatogram alignment, and peak integration from extracted ion chromatograms. Lipid annotation was based on precursor (mass tolerance 4 ppm) and product ions in both positive and negative ion mode. Individual lipids were expressed as ratios to internal standard specific for each lipid class, then multiplied by the amount of internal standard added to produce a molar amount of each lipid per sample, which was normalised to mg extracted tissue.

#### Targeted Lipidomics

sphingolipid quantification was performed by selected reaction monitoring (SRM) on a TSQ Altis triple quadrupole mass spectrometer (Thermo Fisher), operated in positive ion mode. Lipids were separated on a 2.1 × 100 mm Agilent Eclipse Plus C8 column (1.8 μm pore size) with a 24 min chromatography run at a flow rate of 300 μL/min. The HPLC gradient conditions were as follows: 0 min, 20:80 A/B; 2 min, 20:80 A/B; 7 min, 13:87 A/B; 14 min, 0:100 A/B; 20.5 min, 0:100 A/B; 21 min, 20:80 A/B; 24 min, 20:80 A/B. Solvent A: 0.2% formic acid, 2 mM ammonium formate in MilliQ water; Solvent B: 1% formic acid, 1 mM ammonium formate in methanol. Peak integration was carried out using Xcalibur software (Thermo Fisher) and peaks were normalised as ratios to their class specific internal standard (d18:1/12:0 SM, d18:1/12:0 HexCer, d18:1/12:0 sulfatide, d18:1/17:0 ceramide, d17:1 sphingosine and d17:1 S1P). Specific SRM transitions used to identify each lipid are given in the relevant supplementary data tables. The *m*/*z* 264.3 and 262.3 product ions were used to distinguish lipids with a sphingosine or a sphingadiene base, respectively. In the case of SM species, both the dominant *m*/*z* 184.1 product ion and a second qualifying *m*/*z* 264.3 or 262.3 product ion were used for identification. All detected sphingolipids were verified as conforming to our previously identified quadratic elution profile (30). For analysis of sphingosine, sphingadiene, S1P, and sphingadiene 1-phosphate, a previously-described method was used (23), with the inclusion of a *m*/*z* 298.3 → 262.3 transition for sphingadiene and *m*/*z* 378.3 → 262.3 for sphingadiene 1-phosphate.

### Lipid characterisation by 193 nm UVPD-MS/MS

193 nm UVPD-MS/MS was implemented on a custom modified Q-Exactive Plus Orbitrap mass spectrometer (Thermo Fisher), using the output from a Coherent ExciStar 500 XS ArF excimer laser (Santa Clara, CA, USA), as previously described (12). Lipid solutions were prepared by placing 10 μL of lipid extracts into the wells of a PTFE 96-well plate. The samples were dried down in a GeneVac miVac sample concentrator (SP Scientific, Warminster, PA, USA) then resuspended in 40 μL propan-2-ol:methanol:chloroform (4:2:2, v:v:v) containing 2 mM lithium acetate. The plate was then sealed with Teflon Ultra-Thin Sealing Tape (Analytical Sales and Services, Pompton Plains, NJ, USA) prior to introduction to the mass spectrometer via nanoESI (nESI) using an Advion Triversa Nanomate (Ithaca, NY, USA) operating at a spray voltage of 1.3 kV and a gas pressure of 0.3 psi. The ion source interface was set to operate at an inlet temperature of 300 °C and S-Lens value of 50%. All nESI-UVPD-MS/MS spectra were acquired in the Orbitrap mass analyzer using a mass resolving power of 70,000 (at *m*/*z* 400). The AGC target was maintained at 1 × 10^5^ and the maximum injection time was set to 500 ms. Precursor ions were mono-isotopically isolated using an isolation window of ±1.0 *m*/*z*. UVPD-MS/MS spectra were generated using 100 laser pulses/scan (500 Hz repetition rate, approximately 4-5 mJ/pulse, max power 2.5 W), during which the HCD collision energy was set to 2 eV. Spectra shown are the average of 50 scans. The C=C double bond positions in sphingadiene containing lipids were determined based on the diagnostic product ions formed from 193-nm UVPD fragmentation, as described previously (12).

### Western blotting

Cells were lysed in RIPA buffer (20 mM Tris-HCl, pH 7.4, 100 mM NaCl, 1 mM EDTA, 0.1% SDS, 0.5% sodium deoxycholate, 1% Triton X-100, 10% glycerol, 1 mM NaF, 2 mM Na4P2O7, cOmplete™ EDTA-free protease inhibitor cocktail). RIPA buffer extracts (12.5 μg) were resolved on to 4-12% Bolt gels, then transferred to polyvinylidene fluoride membranes and blocked for 1 h with tris buffered saline containing 0.1% Tween-20 (TBST) and 5% skim milk powder. Membranes were washed 3 times with TBST, then cut in half at 50 kDa and incubated overnight rabbit-anti-SphK2 (Cell Signaling Technology, #32346, 1:1000 dilution) or rabbit anti-actin (Abcam, #ab8227, 1:5000 dilution), in TBST with 5% bovine serum albumin. Membranes were washed 3 times with TBST, incubated for 1 h in TBST/5% skim milk with anti-rabbit-HRP (Cell Signaling Technology, #7074, 1:5000 dilution), washed another 3 times, then imaged using chemiluminescence reagent (Merck, #WBKLS0500) and a BioRad ChemiDoc Touch imager. Densitometry was performed using BioRad ImageLab software version 5.2.

### Statistical analysis

For both untargeted and targeted lipidomic analysis, levels of each lipid were compared between SphK2^−/−^ and WT mice (or SphK1^−/−^ and WT) using a two-tailed unpaired t-test. Values were first log-transformed (natural log) to achieve a normal distribution for t-tests. P-values were adjusted for multiple comparisons using the false discovery rate approach of Benjamini, Krieger, and Yekutieli, with Q < 0.1 and 0.05 considered significant for untargeted and targeted LC-MS/MS, respectively (GraphPad PRISM). Complete lists of unadjusted P-values and adjusted Q-values are reported in Supplementary Tables 1-4. For D7-sphingosine pulse-chase, the effect of incubation time and genotype (shCtrl vs shSphK2) on the levels of each lipid were analysed by 2-way ANOVA (GraphPad PRISM). Dunnett’s post-test was applied to test for differences at each time point relative to time = 0.

## Results

### Accumulation of sphingadiene lipids in the hippocampus of SphK2 deficient mice

Untargeted lipidomic (LC-MS/MS) analysis of hippocampus tissue from SphK1^−/−^ and SphK2^−/−^ mice, along with WT control littermates for each, identified phospholipids (phosphatidylcholine, PC; phosphatidylethanolamine, PE; phosphatidylserine, PS), sphingolipids (ceramide, Cer; sulfatide, ST; hexosylceramide, HexCer; sphingomyelin, SM), and neutral (diacylglycerol, DAG; and triacylglycerol, TAG) lipid species. Lipids identified using LipidSearch software were manually verified for peak shape and correct product ions, resulting in a total of 357 confirmed lipids in SphK2^−/−^ and 363 in SphK1^−/−^ mice. After adjusting for false discovery rate, a total of thirteen lipids were significantly increased in SphK2^−/−^ compared to WT mice (Figure 2A and Table 1). In contrast, no lipids differed significantly in abundance between SphK1^−/−^ mice and their WT littermates (Figure 2B). All thirteen lipids whose levels were significantly increased in SphK2^−/−^ mice were sphingolipids containing a sphingadiene (18:2) long chain base, as identified by the diagnostic *m*/*z* 262.252 product ion (expected *m/*z 262.253) that distinguishes sphingadiene from sphingosine (*m*/*z* 264.269) and dihydrosphingosine (*m*/*z* 266.284). CID-MS/MS product ion data for 18:2/18:0 ceramide and SM are illustrated as examples (Figure 2C-D). The C18:0 N-acyl chain is the most abundant fatty acyl chain sphingolipid for both ceramide and SM in the brain. The full dataset is available as Supplementary Table 1 and 2.

**Table 1.** Lipids whose levels are significantly different in hippocampus of SphK2 ^−/−^ compared to WT mice. Both unadjusted P values, and P values adjusted for multiple comparisons (Q values) are reported. MW: molecular weight; Cal. *m*/*z*: calculated *m*/*z*; Obs. *m*/*z*: observed *m*/*z*.

| Lipid | MW | Cal. *m/z* | Obs. *m/z* | Ion | Mean ± SD<br>(nmoles/mg tissue) |  | P value | Q value |
| --- | --- | --- | --- | --- | --- | --- | --- | --- |
|  |  |  |  |  | WT | SphK2<sup>-/-</sup> |  |  |
| Cer(d18:2/18:0) | 563.5277 | 564.5350 | 564.5343 | H+ | 0.943 ± 0.410 | 4.558 ± 1.649 | 0.0001 | 0.0381 |
| CerG1(d18:2/21:0+O) | 783.6224 | 784.6297 | 784.6297 | H+ | 0.010 ± 0.006 | 0.064 ± 0.038 | 0.0003 | 0.0466 |
| Cer(d18:2/22:0) | 619.5903 | 620.5976 | 620.5953 | H+ | 0.005 ± 0.002 | 0.016 ± 0.003 | 0.0004 | 0.0466 |
| Cer(d18:2/22:1) | 617.5747 | 618.5820 | 618.5818 | H+ | 0.042 ± 0.028 | 0.152 ± 0.062 | 0.0011 | 0.0954 |
| SM(d38:2) | 756.6145 | 757.6218 | 757.6210 | H+ | 0.049 ± 0.015 | 0.139 ± 0.060 | 0.0015 | 0.0954 |
| SM(d36:2) | 728.5832 | 729.5905 | 729.5903 | H+ | 2.643 ± 1.259 | 7.277 ± 2.998 | 0.0016 | 0.0954 |
| CerG1(d18:2/18:0) | 725.5805 | 726.5878 | 726.5870 | H+ | 0.028 ± 0.016 | 0.096 ± 0.050 | 0.0020 | 0.0954 |
| Cer(d18:2/24:1) | 645.6060 | 646.6133 | 646.6132 | H+ | 0.008 ± 0.006 | 0.030 ± 0.013 | 0.0021 | 0.0954 |
| CerG1(d18:2/18:0+O) | 741.5755 | 742.5828 | 742.5820 | H+ | 0.031 ± 0.018 | 0.095 ± 0.057 | 0.0025 | 0.0954 |
| CerG1(d18:2/20:0+O) | 769.6068 | 770.6141 | 770.6111 | H+ | 0.045 ± 0.027 | 0.197 ± 0.129 | 0.0025 | 0.0954 |
| Cer(d18:2/23:1) | 631.5898 | 632.5976 | 632.5975 | H+ | 0.018 ± 0.013 | 0.053 ± 0.019 | 0.0029 | 0.0954 |
| Cer(d18:2/22:0+O) | 639.6165 | 640.6238 | 640.6237 | H+ | 0.014 ± 0.010 | 0.042 ± 0.017 | 0.0030 | 0.0954 |
| Cer(d18:2/24:2) | 643.5903 | 644.5976 | 644.5974 | H+ | 0.019 ± 0.012 | 0.052 ± 0.023 | 0.0034 | 0.0999 |

**Figure 2.**
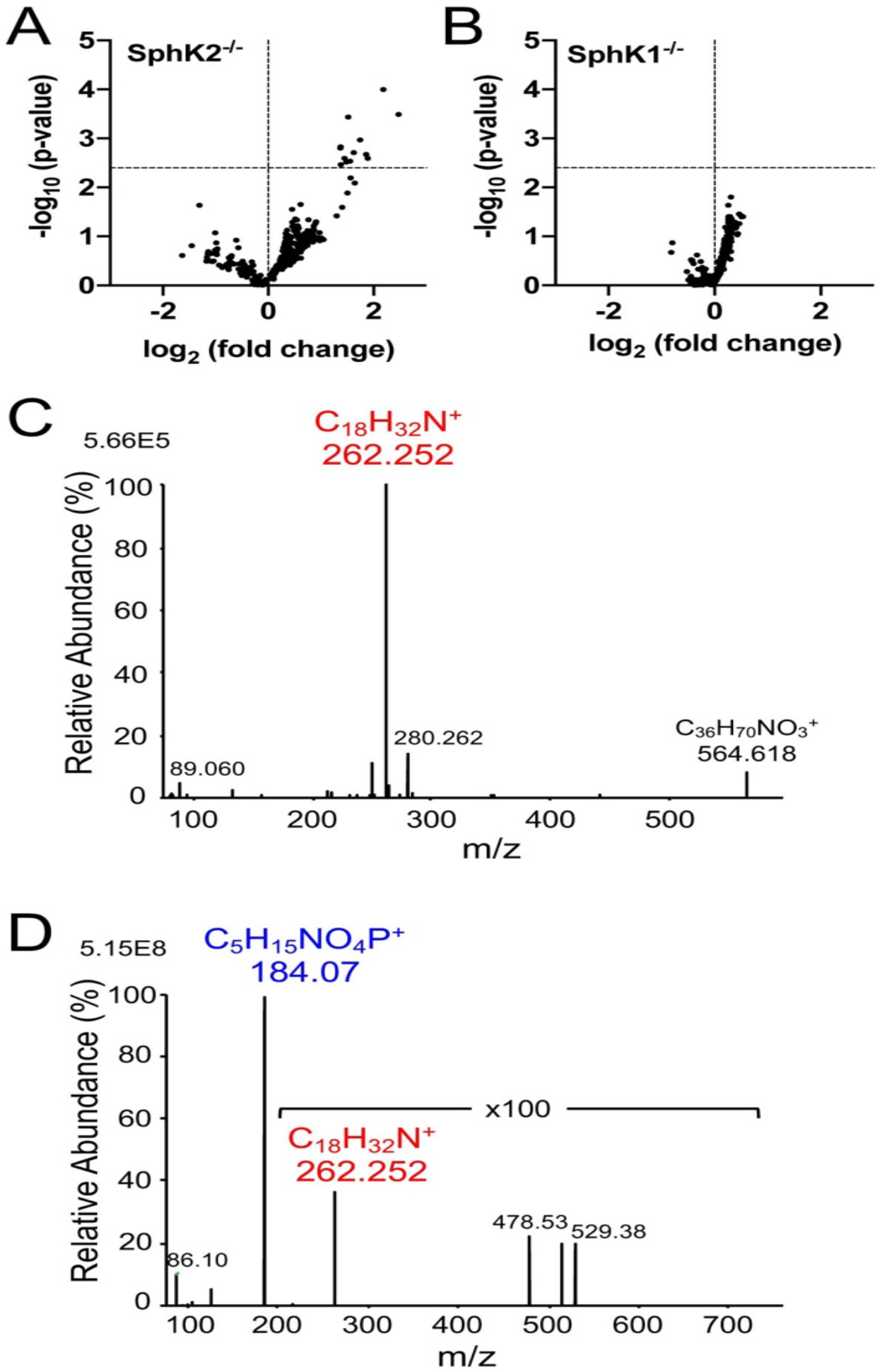
Sphingadiene-containing lipids are increased in hippocampus of SphK2 ^−/−^ mice. (A) Volcano plots showing p-value against fold change for each lipid identified in (A) SphK2^−/−^ and (B) SphK1^−/−^ mice, relative to WT control littermates (mean fold-change from 6 mice per group). Dotted line shows threshold for statistical significance at Q < 0.1. Identities of sphingolipids significantly up-regulated in SphK2^−/−^ mice are given in Table 1. (C-D) Q-Exactive HF-X CID-MS/MS product ion spectra for [M+H]^+^ ions of (C) d18:2/18:0 ceramide and (D) d18:2/18:0 SM. Assignment as d18:2/18:0 ceramide is based on precursor ion *m*/*z* 564.534 and product ion *m*/*z* 262.252, which is diagnostic for sphingadiene. The *m*/*z* 184.073 choline phosphate product ion together with precursor ion *m*/*z* 729.590 are diagnostic of d36:2 SM, and the presence of the *m*/*z* 262.252 product ion allows for the assignment of this lipid as d18:2/18:0 SM.

Our untargeted lipidomic data was validated by targeted LC-MS/MS on hippocampus samples from an independent cohort of 6 month old SphK2^−/−^ mice. Total levels ceramide, SM, HexCer, and sulfatide containing d18:2 sphingadiene backbones were 3.0-, 2.2-, 3.4-, and 2.4-fold higher (all P < 0.0001), respectively, in SphK2^−/−^ compared to WT mice (Figure 3A-D). The increase in sphingadiene lipid levels in SphK2^−/−^ mice was observed for all individual acyl chain variants (Supplementary Table 3). In contrast, the total level of d18:1 ceramides increased by a more modest 1.3-fold (P = 0.021) in SphK2^−/−^ hippocampus samples, and there was no significant difference in total levels of d18:1 SM, HexCer, and sulfatide (Figure 3A-D). Sphingosine and sphingadiene were 2- and 5-fold higher, respectively, in the hippocampus of SphK2^−/−^ mice (Figure 3E, both P < 0.001); whilst S1P and sphingadiene 1-phosphate were 3.5- (P = 0.0002) and 2-fold (P = 0.007) lower, respectively (Figure 3F). The 72% reduction in d18:1 S1P in SphK2^−/−^ mouse brains is in agreement with prior data showing that SphK2 is responsible for the majority of S1P synthesis in the brain (25).

**Figure 3.**
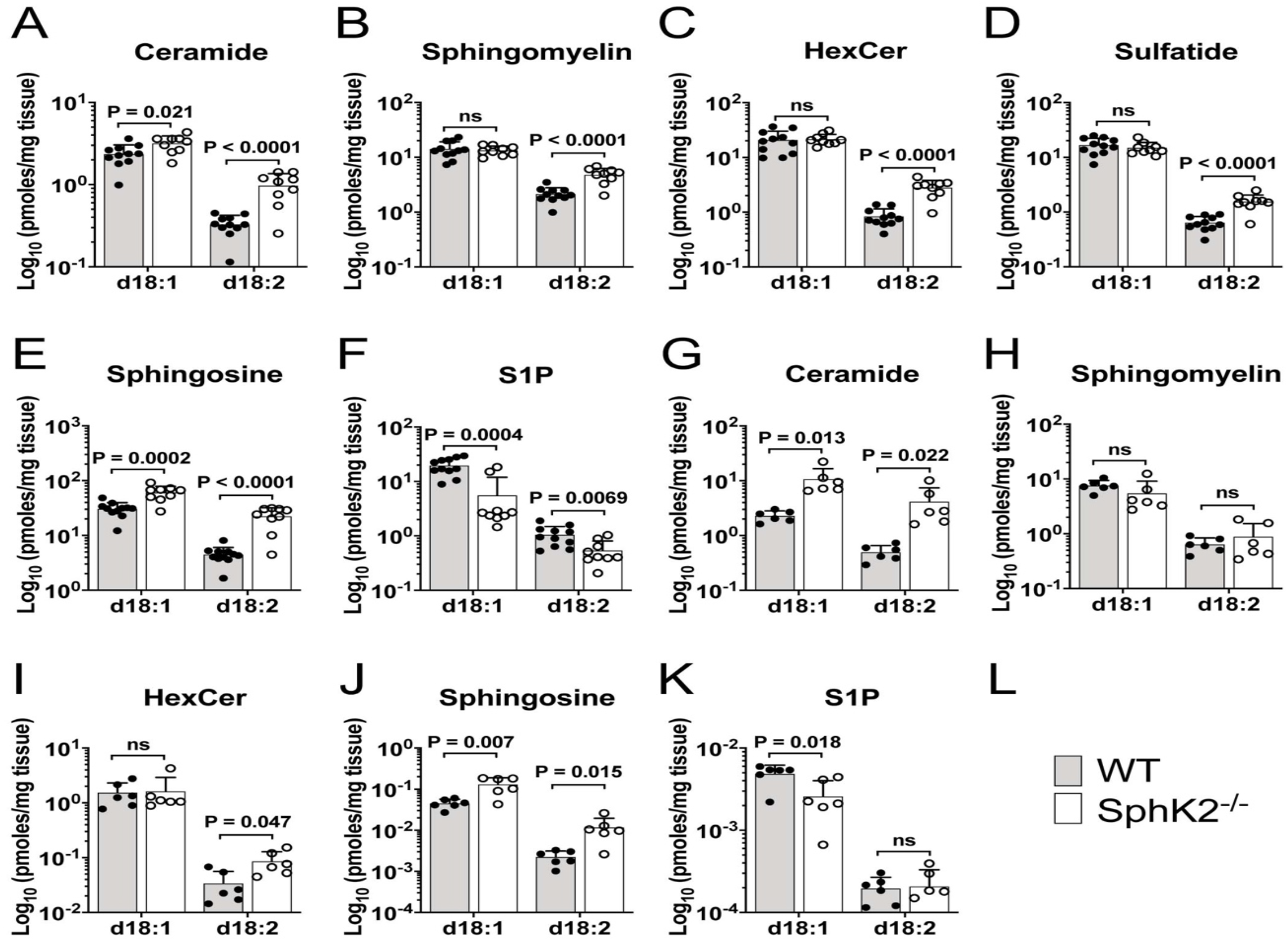
Increased sphingadiene-containing lipids in hippocampus and liver of SphK2 ^−/−^ mice. (A-F) Total levels of d18:1 and d18:2 ceramide (A), SM (B), HexCer (C), sulfatide (D), sphingosine (E) and S1P (F), in hippocampus extracts of WT (shaded bar) and SphK2^−/−^ (clear bar) mice at 6 months of age (n = 11, WT; n = 9, SphK2^−/−^ mice per group), determined by targeted triple quadrupole LC-MS/MS analysis. (G-K) Levels of d18:1 and d18:2 ceramide (G), SM (H), HexCer (I), sphingosine (J), and S1P (K), in livers of WT (shaded bar) and SphK2^−/−^ (clear bar) mice at 8 months of age (n = 6 mice per group). Statistical significance was determined by unpaired t-tests.

To determine if SphK2^−/−^ mice accumulate sphingadiene lipids in another tissue that expresses relatively high levels of SphK2 (31), we also quantified sphingolipids in liver. Ceramides with a d18:1 sphingoid base were 5-fold higher (P = 0.007), whilst those with a d18:2 base were 8-fold higher (P = 0.022) in SphK2^−/−^ compared to WT mouse livers (Figure 3G). SM and HexCer with d18:1 sphingoid bases, as well as d18:2 SM species, did not differ significantly between the two genotypes, whereas d18:2 HexCer was 2.5-fold higher (P = 0.024) in the SphK2^−/−^ livers (Figure 3H-I and Supplementary Table 4). Sphingosine and sphingadiene levels were 3-fold (P = 0.011) and 5-fold (P = 0.0005) higher (Figure 3J), whilst S1P was 2-fold lower (P = 0.018), in SphK2^−/−^ liver tissue (Figure 3K). Sphingadiene 1-phosphate was not significantly altered in SphK2^−/−^ liver tissue. Therefore, as observed in hippocampus, there was a preferential increase of d18:2 relative to d18:1 sphingolipids in livers of SphK2-deficient mice.

### UVPD-MS/MS assigns double bonds to the C4 and C14 position of sphingadiene

To pinpoint the position of the double bonds in the major brain sphingadiene lipids, we employed 193 nm UVPD-MS/MS on a custom modified high resolution Orbitrap instrument. Precursor ions at *m*/*z* 570.542 and 735.597 corresponded to the expected [M + Li]^+^ ions for d18:2/18:0 ceramide and SM, respectively (Figure 4). UVPD-MS/MS of the *m*/*z* 570.542 precursor ion (Figure 4A) yielded product ions at *m*/*z* 243.229 and 332.313, as expected for UV-induced cleavage between C2 and C3 of the sphingadiene base. A pair of product ions differing by 24.000 mass units (*m*/*z* 364.339 and 388.339) were diagnostic for the C4-C5 double bond that is typical of sphingosine. Another pair differing by 24.000 mass units (*m*/*z* 502.480 and 526.480), corresponding to loss of C_5_H_8_ and C_3_H_8_ resulting from cleavage of the C13-C14 and C15-C16 bonds in the long chain base (Figure 4A), confirm the presence of the second unsaturated bond in the sphingoid base chain and localize it to the C14-C15 position. The product ion at *m*/*z* 302.2666, resulting from cleavage of the amide bond and subsequent loss of C_18_H_36_O, confirms the C18:0 saturated acyl-chain amide-linked to sphingadiene. All product ion masses matched the expected masses to within 4 parts per million (ppm).

**Figure 4.**
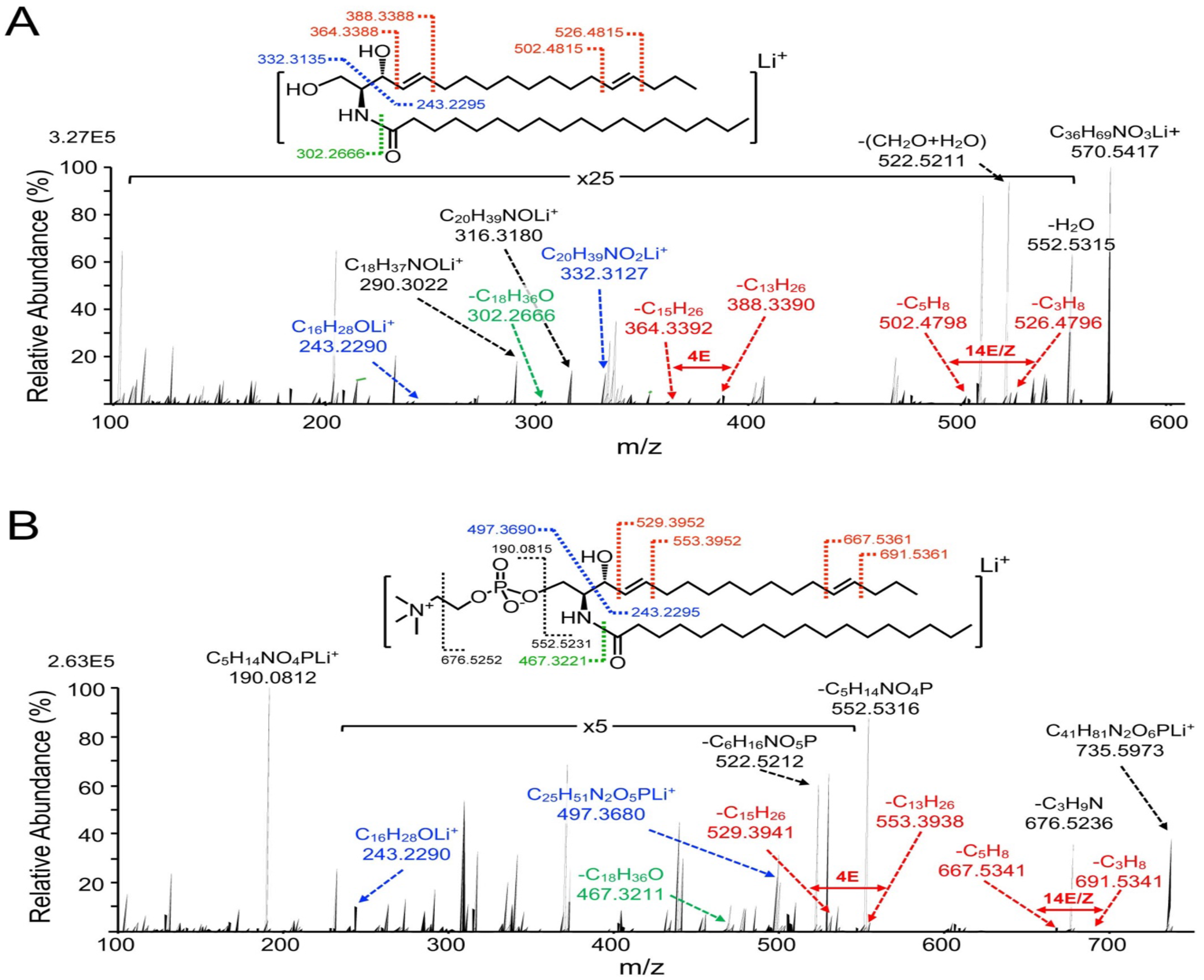
193 nm UVPD-MS/MS spectra for 18:2 ceramide and SM. Product ion spectra resulting from UVPD-MS/MS of [M + Li]^+^ precursor ions of (A) d18:2(4,14)/18:0 ceramide and (B) d18:2(4,14)/18:0 SM. The chemical structure above each spectrum shows points of cleavage, labelled with the expected *m*/*z* for the resulting product ions. Structurally diagnostic product ions for sphingoid backbone (blue), N – C amide bond (green) and specific product ions resulting from cleavage of the sphingoid base double bonds (red) have been highlighted.

UVPD-MS/MS of the *m*/*z* 735.597 [M + Li]^+^ d18:2/18:0 SM ion yielded products at *m*/*z* 243.229 and 497.369, resulting from cleavage between C2 and C3 of the sphingoid base (Figure 4B). Two pairs of product ions differing by 24.000 mass units were also observed from cleavage at the C4-C5 (*m*/*z* 529.394 and 553.394) and C14-C15 positions (*m*/*z* 667.534 and 691.534), indicative of a sphinga-4,14-diene backbone. The C18:0 N-acyl chain was confirmed by the amide bond fragmentation product ion with *m*/*z* 467.322, which corresponds to a loss of C_18_H_36_O. A strong signal was detected for product ion *m*/*z* 190.081, which corresponds to the lithium adduct of the SM choline-phosphate headgroup.

### Both SphK1 and SphK2 exhibit similar affinity for sphingosine and sphinga-4,14-diene

SphK2 is quantitatively more important for S1P synthesis in the brain than SphK1 (Figure 3F) (25), and demonstrates broader substrate specificity, as it is able to catalyse the phosphorylation of phytosphingosine and the sphingoid base analog Fingolimod, as well as sphingosine and dihydrosphingosine (31, 32). As sphingosine kinases catalyse an essential step in the catabolism of all sphingolipids, we assessed whether accumulation of sphingadiene-containing lipids may be attributed to poor affinity of SphK1 for sphingadiene as a substrate. This would result in selective accumulation of sphingadiene lipids in the absence of SphK2. We therefore compared phosphorylation of sphingosine and sphinga-4,14-diene over time in brain lysates of SphK1^−/−^ or SphK2^−/−^ mice. Sphingosine kinase activity in these mice is wholly dependent on SphK2 or SphK1, respectively. Deuterated sphingosine (D7-sphingosine) was used to avoid interference from endogenous sphingosine and S1P present in the brain lysates used as the source of enzyme for the reactions. No deuterated standard was available for sphingadiene, and we therefore used the unlabelled standard. Both SphK1 (SphK2^−/−^ brain extracts) and SphK2 (i.e. SphK1^−/−^ brain extracts) phosphorylated sphingadiene with similar efficiency to sphingosine (Figure 5A-D). The *in vitro* phosphorylation rate was much lower in SphK1^−/−^ than SphK2^−/−^ brain extracts. This is in agreement with a recent publication reporting that SphK1 is much more efficient than SphK2 in the *in vitro* sphingosine kinase activity assay, whereas SphK2 deficiency has a much greater effect than SphK1 deficiency on neuronal S1P levels (33).

**Figure 5.**
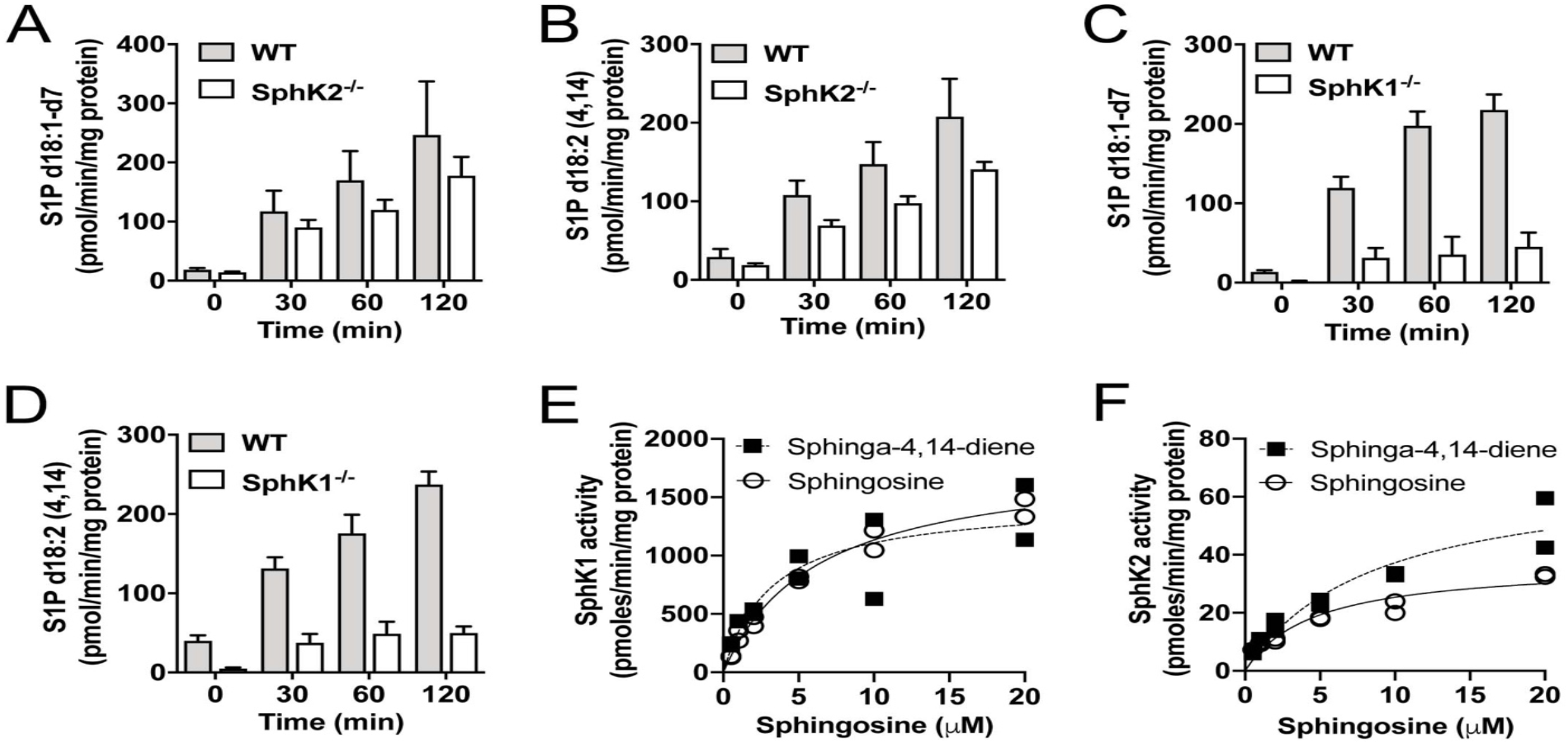
SphK1 and SphK2 show similar affinity for sphingosine and sphingadiene. (A-D) Sphingosine kinase activity of SphK2^−/−^ (A and B) or SphK1^−/−^ (C and D) mouse brain lysates and WT littermate controls, with D7-sphingosine (A and C) or sphinga-4,14-diene (B and D) as substrates. Results show mean and standard error for triplicate assays. (E) SphK1 and (F) SphK2 activity as a function of sphingosine (d18:1; open circles) or sphingadiene (d18:2; closed squares) concentration, measured using recombinant human SphK1 or SphK2. Michaelis-Menten constants (K_m_) and maximal reaction rates (V_max_) were calculated for each substrate using non-linear regression, and are provided in the text.

We also investigated the affinity and maximal reaction rate of recombinant human SphK1 and SphK2 for sphingosine and sphinga-4,14-diene. The Michaelis constant (K_m_) and maximum reaction rate (V_max_) for sphingosine were 6.2 μM and 1832 pmol/min/μg for SphK1; and 4.6 μM and 36.98 pmol/min/μg for SphK2 (Figure 5E and F). These K_m_ values correspond well with previously reported values for SphK1 (5–17 μM) and SphK2 (3-5 μM) (31, 34–36). The K_m_ and V_max_ values for sphinga-4,14-diene were 3.2 μM and 1469 pmol/min/μg for SphK1; and 8.5 μM and 69.10 pmol/min/μg for SphK2 (Figure 5E and F). Therefore, the accumulation of sphingadiene-containing lipids in the absence of SphK2 cannot be explained by poor affinity of SphK1 for sphingadiene.

### Sphingadiene (d18:2) ceramides are metabolised more slowly than d18:1 ceramides

Given that sphingadiene and sphingosine are phosphorylated at equivalent rates by SphK1 and SphK2, we tested if the accumulation of sphingadiene-containing lipids in SphK2^−/−^ brains could be attributed to intrinsically slower turnover of these lipids in comparison to their more abundant d18:1 counterparts. This could cause selective accumulation of sphingadiene-containing lipids in SphK2^−/−^ brains, as the rate of catabolism for all sphingolipids is predicted to be slower in the absence of SphK2. As the majority of brain lipids are synthesized by oligodendrocytes (37, 38), we performed a pulse-chase experiment with the oligodendrocyte cell line MO13.3, first loading cells with deuterated (D7) sphingosine for 2 h, then following the levels of D7-labelled sphingolipids over time after removal of the D7-sphingosine tracer. To test the role of SphK2 in metabolism of labelled sphingolipids, we generated cells stably expressing shRNA against SphK2 (shSphK2) and non-silencing shRNA control cells (shCtrl). Effective silencing of SphK2 with two distinct shRNA sequences (shSphK2-A and shSphK2-B) was confirmed by western blot (Figure 6A).

**Figure 6.**
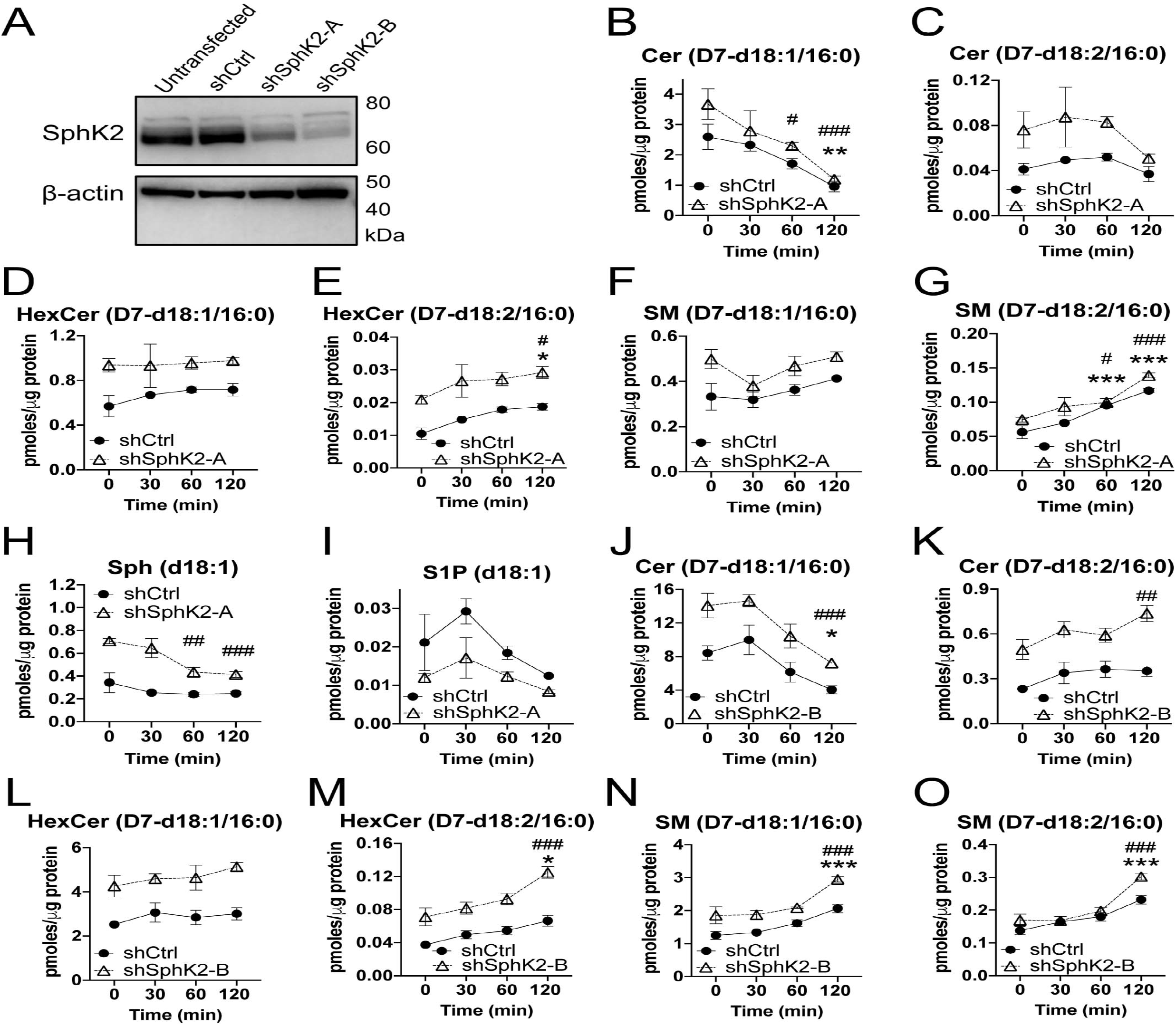
d18:1 ceramides are metabolized more rapidly than d18:2 ceramides. (A) Western blot for SphK2 protein levels in untransfected, shCtrl, shSphK2-A, and shSphK2-B MO3.13 cell pools. (B-G) Levels of D7-d18:1/16:0 (B,D,F) and D7-d18:2/16:0 (C,E,G) ceramide (Cer) (B,C), HexCer (D,E), and SM (F,G) over time, following 2 h loading of shCtrl or shSphK2-A cells with D7-sphingosine (1 μM). (H-I) Levels of D7-sphingosine and D7-S1P. (J-O) Levels of D7-d18:1/16:0 (J,L,N) and D7-d18:2/16:0 (K,M,O) Cer (B,C), HexCer (D,E), and SM (F,G) over time, following 2 h loading of shCtrl or shSphK2-B cells with D7-sphingosine (1 μM). Closed circles (•): shCtrl; open triangles (△): shSphK2. Statistical significance was determined by 2-way ANOVA, and Dunnett’s post-test was applied to compare lipid levels at 30, 60, or 120 min to the 0 min time point. *P<0.05; **P<0.01, ***P<0.001 relative to the 0 time point in shCtrl cells; #P<0.05; ##P<0.01, ###P<0.001 relative to the 0 time point in shSphK2 cells. Overall ANOVA results for shCtrl vs shSphK2 cells are reported in Results.

After loading with D7-sphingosine, we detected quantifiable amounts of D7-labelled sphingadiene-based ceramide, HexCer, and SM. Over 70% were lipids containing a C16:0 N-acyl chain, and we have therefore shown levels of these C16:0 lipids over time (Figure 6B-O). Levels of D7-d18:1/16:0 ceramide, SM, and HexCer synthesized from D7-sphingosine were much higher than their D7-18:2 equivalents in both shCtrl and shSphK2-A cells, whilst both D7-d18:1 and D7-d18:2 sphingolipids were higher in shSphK2-A cells compared to shCtrl cells (p < 0.001 by 2-way ANOVA for d18:1/16:0 ceramide, d18:2/16:0 ceramide, d18:1/16:0 HexCer, d18:2/16:0 HexCer, and d18:1/16:0 SM; P = 0.0025 for d18:2/16:0 SM). Levels of D7-sphingosine remained relatively constant in the shCtrl cells but declined significantly at 60 and 120 min in shSphK2-A cells (Figure 6H), presumably as the sphingosine was converted to other sphingolipids. D7-sphingadiene was not detected, suggesting that the Δ14 desaturase that produces sphinga-4,14-diene acts on ceramide, rather than sphingosine, as its substrate. As expected, D7-S1P was higher in shCtrl compared to shSphK2-A cells (Figure 6I; P<0.0001 by 2-way ANOVA). This may explain the higher levels of D7-ceramide, D7-SM, and D7-HexCer in shSphK2 cells, as flux through SphK2 was inhibited.

D7-d18:1/16:0 ceramide declined 2.7-fold in shCtrl (P = 0.008), and 3.1-fold in shSphK2-A cells (P = 0.0001), 2 h after removal of the D7-sphingosine tracer (Figure 6B). In contrast, levels of D7-d18:2/16:0 ceramide did not decline significantly over the 2 h time course (Figure 6C). Levels of D7-d18:1/16:0 HexCer and D7-d18:1/16:0 SM were not significantly altered over the 2 h time course (Figure 6D and F), whereas both D7-d18:2/16:0 HexCer and D7-d18:2/16:0 SM increased significantly in shCtrl and shSphK2-A cells (Figure 6E and G). Similar data was observed upon repeating the experiment with cells expressing a different shRNA sequence, shSphK2-B (Figure 6J-O). D7-d18:1/16:0 ceramide declined in both shCtrl and shSphK2-B cells (P = 0.032 in the WT cells, P = 0.0008 in shSphK2), whereas D7-d18:2/16:0 ceramide increased 1.5-fold in both shSphK2-B cells and shCtrl cells (P = 0.009 in shSphK2-D; not significant, P = 0.28, in shCtrl), 2 h after removing the D7-sphingosine tracer (Figure 6K). D7-d18:1/16:0 HexCer levels did not change significantly over the 2 h time-course, whereas D7-d18:2/16:0 HexCer increased significantly in both cell lines (Figure 6L-M). In this experiment, both D7-d18:1/16:0 and D7-d18:2/16:0 SM increased significantly over the 2h time-course (Figure 6N-O). Overall, these results demonstrate that d18:1 ceramides are more rapidly metabolised than d18:2 ceramides.

## Discussion

Sphingosine kinases fulfill dual biochemical functions in mammalian biology. Firstly, they produce the physiologically-essential signalling molecule S1P, which signals both through a family of five G-protein coupled receptors, and through intracellular binding targets. Secondly, they catalyse the penultimate step in the catabolism of sphingolipids, as phosphorylation of sphingoid bases is necessary for their degradation by S1P lyase (1, 13, 15.). Here, we establish that sphingolipids with a sphingadiene long chain base accumulate in the hippocampus of SphK2^−/−^ but not SphK1^−/−^ mice. Using the emerging fragmentation method of UVPD-MS/MS, we localized the double bonds to the C4-C5 and C14-C15 positions of the sphingoid base chain, allowing us to test the rate of phosphorylation of this sphingoid base by SphK1 and SphK2 *in vitro*. There was no intrinsic difference in the capacity for either SphK1 or SphK2 to phosphorylate sphinga-4,14-diene compared to sphingosine. Instead, we show that d18:1 ceramide levels decline over a 2 h time-course in cultured cells, whereas d18:2 ceramide levels do not. Whereas d18:1 HexCer levels were relatively unchanged over the 2 h time course, d18:2 HexCer levels increased. These results suggest that ceramides with a sphinga-4,14-diene backbone are less efficiently catabolized than the more common sphingosine-based lipids. As SphK2 catalyses the bulk of S1P synthesis in the brain (24, 25, 33), the less efficient catabolism of sphingadiene lipids combined with SphK2 deficiency results in abnormal accumulation of these lipids.

Our findings are strongly supported by a very recent publication showing that, compared to S1P, sphinga-4,14-diene 1-phosphate is less efficiently catabolized by S1P lyase *in vitro* (using cell extracts) (39). Our data extends this work by demonstrating that sphingadiene-containing lipids are more slowly catabolized than sphingosine-containing lipids in living cells. Similarly, S1P lyase mutants in the fly *Drosophila melanogaster* accumulate sphinga-4,6-diene (54), which is speculated to contribute to muscle degeneration (40). In agreement with our data, Jojima et al (39) also found that both SphK1 and SphK2 catalyse sphingadiene phosphorylation as efficiently as sphingosine. We note that in their study, Jojima et al did not use a mass spectrometry approach to specifically resolve double bond positions in sphingadiene, instead using synthetic sphinga-4,14-diene lipid standards in combination with HPLC elution time to indicate the double bond position in biological extracts.

In our study, the recently developed experimental technique of UVPD-MS/MS was applied to empirically determine the position of the two double bonds in the sphingadienes identified using untargeted lipidomic profiling. Techniques for assignment of double bond position in lipids without chemical derivatization have only recently been developed and are not yet used in routine LC-MS/MS analyses. UVPD-MS/MS has been used for isomer differentiation and localisation of double bonds in glycerophospholipids (8), lipooligosaccharides (9, 11), gangliosides (10) and sphingolipids (12). However, this is the first report employing UVPD-MS/MS in combination with genetic models to answer specific biological questions concerning lipid metabolism and function. UVPD-MS/MS does not require any derivatisation, modification, or chromatographic separation to resolve sites of unsaturation, making this a powerful technique that can be readily incorporated into lipidomic workflows. An alternative MS/MS approach for double bond assignment is ozonolysis, which coverts alkene bonds to ozonoids that fragment into product ions pairs diagnostic of double bond position (41, 42). However, an ozone source may create unwanted hazards, the ionisation efficiency of ozonolysis is reportedly reduced in positive ion mode, and the high capillary voltage reduces sensitivity (43). Comprehensive structural characterization of lipids can also be performed using electron induced dissociation (EID) in combination with differential ion mobility spectrometry [also referred to as electron-impact excitation of ions from organics (EIEIO)] (44, 45). Fragment spectra from EID can identify lipid class, number of double bonds, and regio-isomers, although the precise position(s) of double bonds can be difficult to isolate using EID, since single bond fragments interfere with double bond diagnostic peaks (44). An important caveat regarding the general applicability of UVPD-MS/MS and other complex dissociation techniques described above is the complexity of data analysis. Even with CID, lipidomic analysis is very complex and requires both training and significant knowledge of lipid structures. Analysing fragments related to double bond positions increases this complexity and may be challenging to implement into broad lipidomic profiling experiments.

Our assignment of the second double bond in murine brain sphingadiene to the C14-C15 position is in agreement with three publications from the late 1960s in which chemical derivatisation techniques coupled to thin-layer or gas chromatography were used to determine double bond positions in d18:2 SM species of human plasma and aorta samples (46–48). Despite this, the levels and functional significance of sphingadiene-containing lipids has been very sparsely investigated relative to the sphingoid bases sphingosine and sphinganine (dihydrosphingosine), and they are often not included in routine lipidomic analyses. It was very recently shown that d18:2 is the second most abundant sphingoid base in human plasma (49), and that levels of sphinga-4,14-diene in mice are highest in kidney, followed by brain, lung, and then colon (39). Sphingadienes with double bonds in the C4-C5 and C8-C9 positions of the sphingoid base chain are common in plants and fungi such as soybean, corn, wheat and yeast (50, 51). Naturally occurring sphingadienes inhibit both chemically- and genetically-induced colon cancer development in mice (51–53), associated with disrupted Akt membrane translocation and signaling (53), and reduced Wnt transcriptional activity (52). Sphingadienes have also been shown to suppress the growth of neuroblastoma xenograft tumors (54).

Introduction of the C4-C5 trans double bond into ceramide is catalysed by ∆4-dihydroceramide desaturase (DEGS1) (55). Two research teams very recently identified FADS3 as the desaturase catalyzing the addition of the C14-C15 double bond in the long chain base (39, 49). Jojima et al (39) presented evidence indicating that the second double bond in the sphingadiene base is introduced into ceramide, not sphingosine. Our data also suggested that desaturation at the C14-C15 bond occurs in ceramide, not sphingosine, as D7-sphingadiene was below the limit of detection in cell culture, despite abundant levels of D7-sphingosine and clearly detectible D7-d18:2 sphingolipids. In contrast, Karsai et al (49) showed that FADS3 can directly desaturate sphingosine *in vitro*, however inhibiting ceramide synthesis with Fumonisin B1 greatly reduced sphingadiene formation in living cells. Mice lacking FADS3 have reduced brain docosahexaenoic acid levels but were reported to display no major phenotypic abnormalities (56). However the International Mouse Phenotyping Consortium (https://www.mousephenotype.org) (57) report impaired auditory response, decreased circulating phosphates, and increased serum albumin in FADS3 knockout mice.

The C14-C15 double bond in sphinga-4,14-diene is a cis bond (14), which introduces a kink in the acyl chain and is thereby expected to decrease the packing density of the lipid bilayer. Accordingly, Jojima et al (39) showed that a lower proportion of d18:2 HexCer localizes to detergent-resistant membrane domains compared to d18:1 HexCer. The brain is enriched with sphingolipids, so an increased proportion of d18:2(4,14) sphingolipids may have important consequences for neuronal lipid domains and myelin integrity. We have reported that SphK2 deficiency synergises with an amyloid-producing transgene to create severe myelin deficits (24). It is possible that myelin structure and function is destabilized by an increased abundance of sphingadiene lipids in SphK2^−/−^ mice, and this will be investigated in future studies.

In conclusion, we have identified significant accumulation of sphingolipids containing a sphingadiene base in the hippocampus of mice lacking functional SphK2 and have employed the emerging technique of UVPD-MS/MS to assign double bond positions to the C4-C5 and C14-C15 positions. To the best of our knowledge, this is the first study to apply UVPD-MS/MS towards an improved understanding of metabolic pathways and enzyme function. The biology of sphingadiene-containing lipids has not been elucidated, and these lipids have not commonly been assayed in targeted lipidomic analyses. Sphingosine kinases remain targets of interest for cancer therapy, and inhibitors of both SphK1 and SphK2 have been investigated in clinical trials. Our findings demonstrate the importance of analyzing and better understanding the functional significance of sphingadiene lipids, particularly given that circulating levels of these lipids were reported as significantly higher in females (49), and knockout of SphK2, or Bax and Bak (58), has now been shown to alter the balance between sphingadiene- and sphingosine-containing lipids.

## Abbreviations

CID: collision induced dissociation
HexCer: hexosylceramide
LC-MS/MS: liquid chromatography-tandem mass spectrometry
SM: sphingomyelin
SphK1: sphingosine kinase 1
SphK2: sphingosine kinase 2
UVPD: ultraviolet photodissociation

## Acknowledgements

This research was supported by National Health and Medical Research Council (NHMRC)-Australian Research Council (ARC) dementia research fellowship APP1110400 (T.A.C.), NHMRC project grants APP1100626 (A.S.D.) and APP1156778 (G.E.R.), and ARC project grant DP190102464 (G.E.R.). We gratefully acknowledge subsidized access to the Sydney Mass Spectrometry Core Facility.

